# The N34S mutation of SPINK1 may impact the kinetics of trypsinogen activation to cause early trypsin release in the pancreas

**DOI:** 10.1101/2020.08.21.262162

**Authors:** Zhuyezi Sun, István Kolossváry, Dima Kozakov, Miklós Sahin-Tóth, Sandor Vajda

## Abstract

The N34S variant of the trypsin inhibitor SPINK1 is the clinically most significant risk factor for chronic pancreatitis, but the underlying molecular mechanism could not be identified. Molecular dynamics simulations and docking of the generated conformational ensemble of SPINK1 to trypsin show that the mutation reduces the fraction of conformations that can directly participate in productive association, thereby reducing the association rate. The small change is difficult to detect by measuring the kinetics of SPINK1 binding to trypsin. However, kinetic modeling reveals that even a small change in the inhibition rate affects the trypsinogen to trypsin conversion rate at the early stage of the reaction when the trypsin concentration is very low, and the impact is substantially amplified by the autocatalytic mechanism of the conversion. Thus, the slightly reduced inhibition rate shortens the delay in the activation of trypsin release, which is therefore occurs within the pancreas.

## Introduction

It is now firmly established that there is a trypsin-dependent pathological pathway in chronic pancreatitis as accumulation of trypsin in pancreas can lead to autodigestion and tissue injury (Hegyi & Sahin-Toth, 2017). The serine protease inhibitor Kazal type 1 (SPINK1, also known as pancreatic secretory trypsin inhibitor, PSTI) is a 6.2-kDa protein secreted by the pancreatic acinar cells that can potently inhibit trypsin. A large number of studies established that the p.N34S variant of SPINK1 is the clinically most significant risk factor for chronic pancreatitis (Hegyi & Sahin-Toth, 2017; Kereszturi, Kiraly, & Sahin-Toth, 2009; Kiraly, Wartmann, & Sahin-Toth, 2007; Koziel, Gluszek, Kowalik, Chlopek, & Pieciak, 2015; Liddle, 2006; Masson, Chen, Audrezet, Cooper, & Ferec, 2013; Rosendahl et al., 2013; Wang et al., 2013; Zou et al., 2018). According to a meta-analysis, the average carrier frequency is 9.7% in patients and 1% in controls and the average odds ratio (OR) is 11(Aoun et al., 2008). Despite efforts to uncover the disease-relevant functional effect of the p.N34S variant, the underlying molecular mechanism is yet to be identified (A Boulling, Chen, Callebaut, & Férec, 2012; Chen & Ferec, 2009; Pfutzer et al., 2000). Witt et al.(Witt et al., 2000) and Pfützer et al.(Pfutzer et al., 2000) hypothesized that the variant causes impaired trypsin inhibition, which would result in elevated intrapancreatic trypsin activity and pancreatic injury. However, experimental studies failed to demonstrate that the p.N34S variant has any effect on the trypsin inhibitory activity or the cellular folding and secretion of SPINK1 (A Boulling et al., 2012; A. Boulling et al., 2007; Kiraly et al., 2007; Kuwata et al., 2002). Attempts to use differences between the structures of unbound SPINK1 and the structure of the inhibitor bound to chymotrypsin to explain the impact of the mutation are far from convincing(A Boulling et al., 2012). A functional role for the co-segregating four intronic alterations has been also excluded (Kereszturi et al., 2009; Masamune et al., 2007).

Here we show that understanding the impact of the SPINK1 mutation it is necessary to consider the activation of trypsinogen by trypsin, which is affected by SPINK1, rather than only the isolated inhibition process. We use molecular dynamics (MD) simulations of the wild-type and the N34S SPINK1, protein-protein docking of the conformational ensembles, and kinetic modeling of trypsinogen activation in the presence of SPINK1 to provide a plausible explanation why the mutation leads to chronic pancreatitis in most patients. The residue N34 of SPINK1 does not face the active site of trypsin, but it is located in a loop region that substantially interacts with the enzyme. As expected for a small protein, the loop in question is very flexible and even in the short time scale of the MD simulations samples a large conformational ensemble. It is well known in protein-protein association kinetics that only some fraction of these conformations lead to productive collisions that yield stable association of the two molecules (Berg & von Hippel, 1985; Camacho, Weng, Vajda, & DeLisi, 1999; Kozakov et al., 2014; von Hippel & Berg, 1989b). Our key finding, based on the MD simulations and protein-protein docking, is that the N34S mutation reduces the flexibility of the loop and the fraction of such productive conformations, thereby reducing the rate of trypsin inhibition by the N34S variant of SPINK1. Although the change in the association rate is small, modeling the kinetics of the trypsinogen to trypsin conversion in the presence of SPINK1 reveals that even the small reduction can have a large effect on the rate of trypsinogen activation, leading to early release of trypsin in the pancreas. The key factors causing such a large impact are the small initial concentration of trypsin and the autocatalytic mechanism of trypsinogen activation that makes the rate of trypsin release highly sensitive to that early trypsin concentration. The importance of the rate of trypsinogen autoactivation in trypsin release has been previously validated (Geisz & Sahin-Toth, 2018). Since SPINK1 binding to trypsin has both high affinity and high association rate, we will demonstrate that the small change in the latter is difficult to measure in experiments. In fact, the reported values for the association rate vary from 1.1×10^6^ M^−1^s^−1^ (Kuwata et al., 2002; Vincent & Lazdunski, 1972) to 3.0×10^6^ M^−1^s. This suggests that the best way to validate the mechanism proposed here would be to measure the rate of trypsinogen activation at low initial trypsin concentrations in the presence of wild-type or N34S inhibitor. However, due to its intrinsic zymogen activity, trypsinogen always includes trace amounts of trypsin, which reduces the reproducibility of results in this type of experiments and emphasizes the need for a computational approach.

## Results

The analysis proceeds in three largely independent steps involving different methodologies. First we use molecular dynamics (MD) to generate conformational ensembles for both the wildtype and the N34S mutant of SPINK1 in order to examine the change in loop dynamics introduced by the mutation. Second, after clustering the frames from the simulations, a number of representative structures from the MD generated conformational ensembles of the two SPINK1 variants are docked to human trypsin to explore the interaction (Kozakov, Brenke, Comeau, & Vajda, 2006). The docking yields a large number of protein complexes, a fraction of which are close to the native structure. Such docked poses represent encounter complexes that are potential transient states along the protein association pathway (Kozakov et al., 2014) and represent productive collisions that will result in inhibition of trypsin. According to MD and docking simulations, the N34S mutation yields changes in the distribution of SPINK1 conformations that reduce the fraction of productive encounters. We will argue that this results in a 10% to 20% reduction in the trypsin-SPINK1 association rate. In the third step of our analysis, we construct a simple kinetic model to demonstrate that, due to the autocatalytic mechanism of the trypsinogen to trypsin transition, even the relatively small reduction in the association rate substantially affects the kinetics of inhibition and results in early release of trypsin in the pancreas.

### Protein structures

Two structures of the human SPINK1 are available in the Protein Data Bank (PDB), one separately crystallized (PDB ID 1hpt) (Hecht, Szardenings, Collins, & Schomburg, 1992), and the second co-crystallized with chymotrypsinogen (PDB ID 1cgi) (Hecht, Szardenings, Collins, & Schomburg, 1991). In the separately crystallized form, the N34 residue in the structure adopts an unusual high-energy conformation in the left-handed alpha-helical region of the Ramachandran plot (A Boulling et al., 2012). Although such conformation may be possible for asparagine (Deane, Allen, Taylor, & Blundell, 1999), we determined it to be an artifact from crystal contact. Running simulations with crystal contact may lead to structures being trapped in a pre-fixed stable conformation, and thus it was undesirable to start MD with the PDB structure 1hpt. More importantly, it was beneficial to start the MD simulations from a conformation closest to the expected bound form, so we decided to focus on the bound conformation of human SPINK1 as the starting structure of our simulations. The inhibitor cocrystallized with chymotrypsinogen in the PDB structure 1cgi is a variant of the human SPINK1 that includes four mutations: K41Y, I42E, D44R and N52D, and we mutated these four residues back to the original SPINK1 sequence. Representative structures from the SPINK1 ensemble generated by molecular dynamics were docked to the structure of human trypsin extracted from the human trypsin-bovine pancreatic trypsin inhibitor (BPTI) complex (PDB ID 2ra3) (Salameh, Soares, Hockla, & Radisky, 2008). The trypsin structure in this complex contains two mutations (Salameh et al., 2008), S195A and R117H (conventional chymotrypsin numbering), that allowed for improved crystallization results. For more accurate docking results, we mutated A195 and H117 back to match the original trypsin sequence.

Since there is no human SPINK1 - human trypsin structure in the PDSB, we developed a model of the complex to be used as the reference native state. To model the native interface, we superimposed the already mentioned human trypsin-bovine pancreatic trypsin inhibitor (BPTI) complex (PDB ID 2ra3) (Salameh et al., 2008) onto the chymotrypsinogen-human SPINK1 complex (PDB ID 1cgi). Chymotrypsinogen is the inactive precursor of chymotrypsin, and is converted into fully active enzyme through the cleavage of a single peptide bond. Chymotrypsin and trypsin share high sequence identity and also very similar tertiary structures (Ma, Tang, & Lai, 2005). In fact, the backbone of the key interactions at the interface loops are nearly identical, and hence chymotrypsin is a good template to model trypsin. The modeled interface of the human trypsin-human SPINK1 complex then became the reference structure later used for evaluating docked near-native poses.

### Molecular dynamics simulations of wild-type and N34S SPINK1

We performed MD simulations both on the wild-type SPINK1 structure extracted from the complex co-crystalized with chymotrypsinogen (PDB ID 1cgi) and on the N34S mutant. Each simulation started with an equilibration and relaxation protocol to remove clashes, and was run for 500 ns with recording intervals of 1000 ps. For each protein, we performed 10 independent simulations, each starting with randomized initial atomic velocities. The resulting 10 trajectories were concatenated to represent an aggregate of 5 μs MD simulation. As shown in **Figures 1a and 1b**, both structures tend to move away from the bound conformation by a few angstroms. The RMSD values were calculated referencing the bound conformation of SPINK1 in the PDB structure 1cgi based on all Ca atoms. In terms of this overall RMSD the N34S mutation slightly reduces the spread of the structures, and moves the maximum of the distribution closer to 2 Å. The two distributions are similar, but the two-sample Kolmogorov-Smirnov test rejects the null hypothesis that the two sets of data are from the same distribution (p < 0.05), and more detailed analysis shows differences in local flexibility.

**Figure 1.**
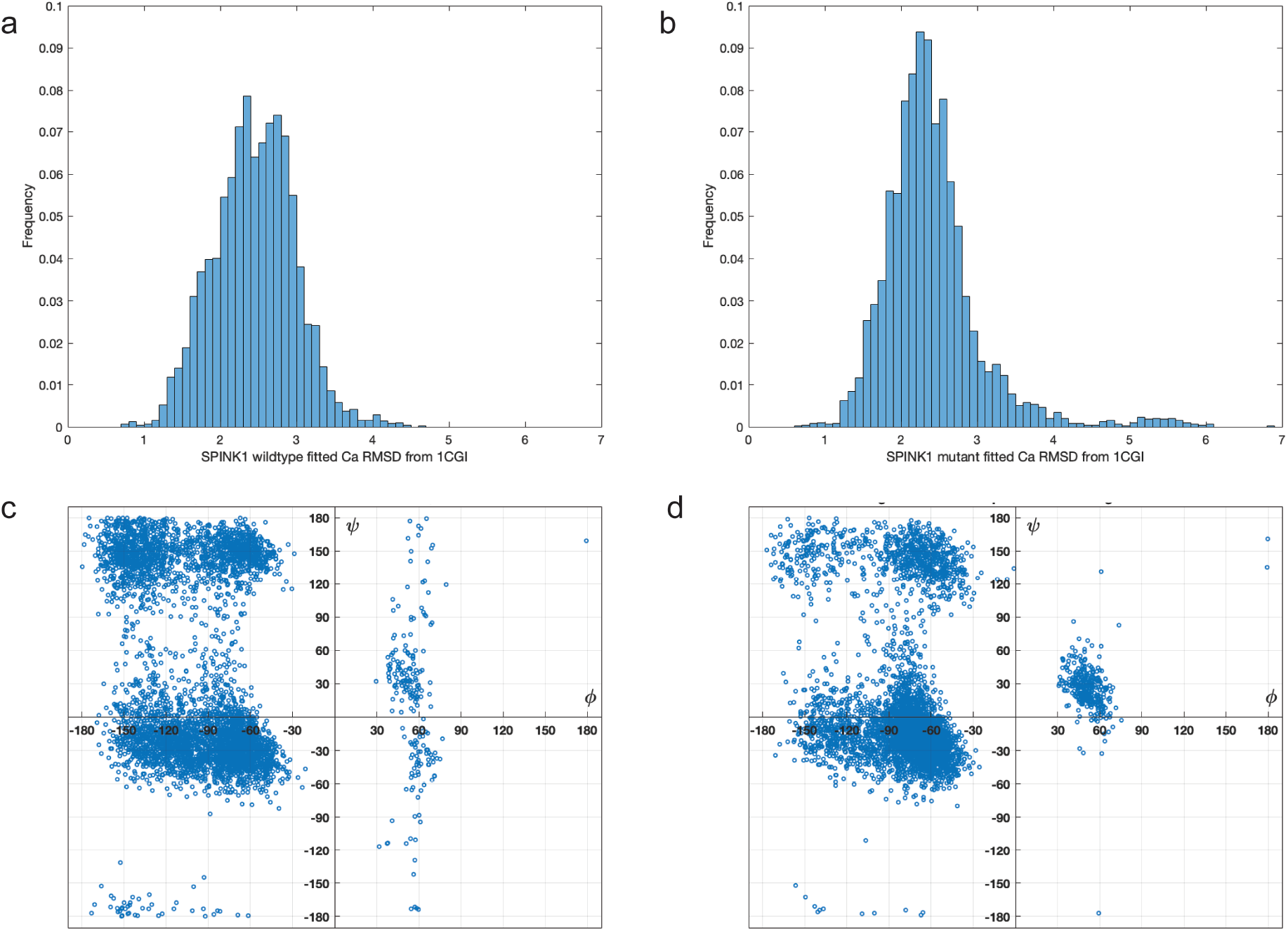
Interface Root Mean Square Deviation (**I**RMSD) histograms and Ramachandran plots for ensembles of structures generated by molecular dynamics (MD) for the wild-type and N34S mutant of SPINK1. **a**. IRMSD values between the SPINK1 structures generated by molecular dynamics simulations of the wild-type SPINK1 and the native structure of the trypsin-bound SPINK1, based on the PDB structure 1cgi. **b**. Same as Figure **1a** for the SPINK1 N34S variant. **c**. Dihedral angles φ and ψ of the residue at position N34 of the MD generated structures for the wild-type SPINK1. **d**. Dihedral angles φ and ψ of the residue S34 of the MD generated structures for the N34S SPINK1.

Although the loop region that includes N34 is stabilized by disulfide bridges between C32-C61 and C39-C58, it is the most flexible part of the protein. **Figures 1c and 1d** shows the backbone dihedral angles φ and ψ of the residue at position 34 of SPINK1 on the Ramachandran plot. The large cluster around (−150°, +150°) in the wild-type protein becomes substantially smaller in the mutant, which results in a less variable loop conformation. In addition, in the wild-type the points on the right side of the Ramachandran plot broadly distribute over the entire −180° < ψ <180° range (Figure 1c), whereas in the mutant are concentrated in the −10° < ψ < 60° region (Figure 1d), also indicating reduced flexibility. The two-sided Wilcoxon rank sum test applied to φ and ψ values of the conformational ensembles generated by MD rejects the null hypothesis of equal medians at the default 5% significance level in both cases, confirming that the differences in the distributions of dihedral angles are significant. To demonstrate the conformational changes of the loop containing residue 34 during the MD simulations of SPINK1, **Figures 2a**, **2b, 2c**, and **2d** show trypsin as grey surface, the trypsin-bound SPINK1 from the structure 1cgi as red cartoon, and a few randomly selected frames from the simulation in various colors. In Figure 2a we face the active site of the enzyme. This view demonstrates that in the trypsin-SPINK1 complex the loop of residues 33 to 39 of the inhibitor fits into an elongated surface cavity of trypsin defined by L97-T98-I99 (on the left from the loop in Figure 2a) and G216-S217-S218 on the right. Superimposing the MD frames on the trypsin structure shows that the loop predominantly moves in the direction away from the enzyme (shown as downward extension in Figure 2a), and clashes with it only in some of the conformations. In contrast, the same loop In the N34S mutant becomes slightly more rigid, its outward motion is more limited, and a higher fraction of frames would clash with the enzyme (Figures 2c and 2d). In particular, residues N37 and G39 of the mutant SPINK1 tend to clash with the side chain of W215 of trypsin. It is well understood that many nonsynonymous single nucleotide polymorphisms are disease causing due to effects at protein-protein interfaces(David, Razali, Wass, & Sternberg, 2012). The disruption of the protein-protein interaction usually occurs due to the loss of an electrostatic salt bridge, the reduction of the hydrophobic effect, the formation of a steric clash, or the introduction of a proline altering the main-chain conformation(David et al., 2012). What is unusual about the N34S mutation of SPINK1 that while N34 is in the interface, its side chain orients away from trypsin, and hence the mutation leads to more subtle effects by changing the flexibility of the loop. However, the role of flexibility in affecting association rates(Greives & Zhou, 2012; Zhou & Bates, 2013) and as a determinant of functional significance of missense mutations(Ponzoni & Bahar, 2018) has been shown for several proteins.

**Figure 2.**
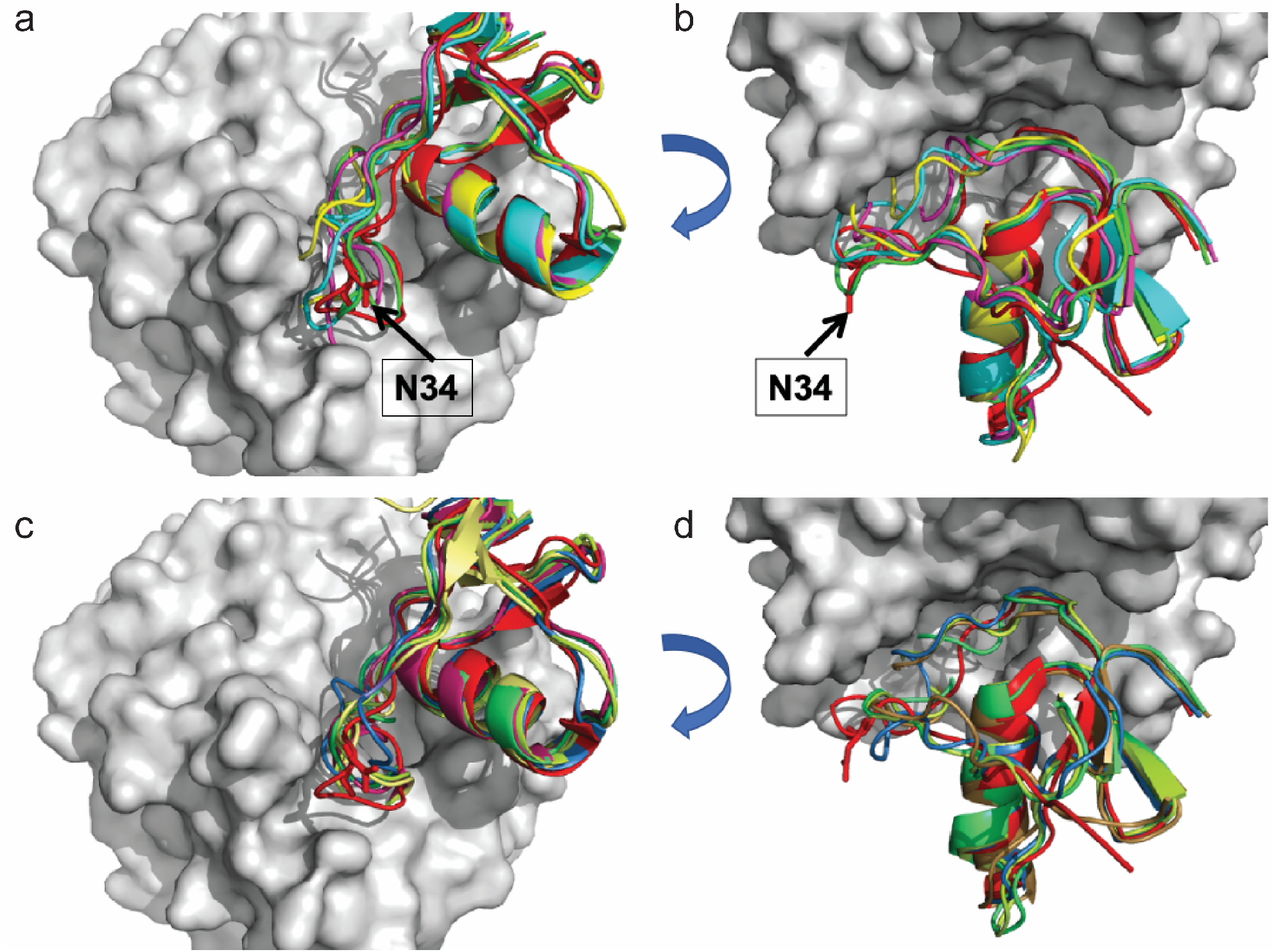
Frames from the MD simulation of wild-type and N34S SPINK1. The trypsin structure (grey surface) and the bound SPINK1 (red cartoon) are superimposed from the 1cgi structure for reference. **a**. Simulation of the wild-type SPINK1 in a view facing the active site of trypsin. **b**. Side view of the wild-type SPINK1 structures shown in Figure 2a. **c**. Simulations of the N34S variant of SPINK1 in the same view as in Figure 2a. **d**. Side view of the N34S variant of SPINK1 structures shown in Figure 2c. Some of the structures clash with trypsin and disappear in the grey surface of the protein.

To extract representative structures from the MD simulations, we subjected the trajectories to clustering. The clustering radii were tailored using the pairwise RMSD distributions as based on our previous experience (Ignatov et al., 2019; Kozakov, Clodfelter, Vajda, & Camacho, 2005), resulting in 2.8 Å and 2.6 Å for the wild-type and the N34S mutant, respectively. Finally, the cluster centers, which were structures with the largest numbers of neighbors, were extracted from the trajectories. Clusters with less than 2% of the total population were removed. Based on these criteria, 12 wild-type cluster centers and ten N34S mutant cluster centers were extracted as representative structures from the simulations. The probability *p_i_* of the ith cluster was defined as = *N_i_/N*, where *N_i_* Is the number of frames in the ith cluster and *N* = ∑*N_i_* is the total number of frames in all retained clusters (**Figure 3a**).

**Figure 3.**
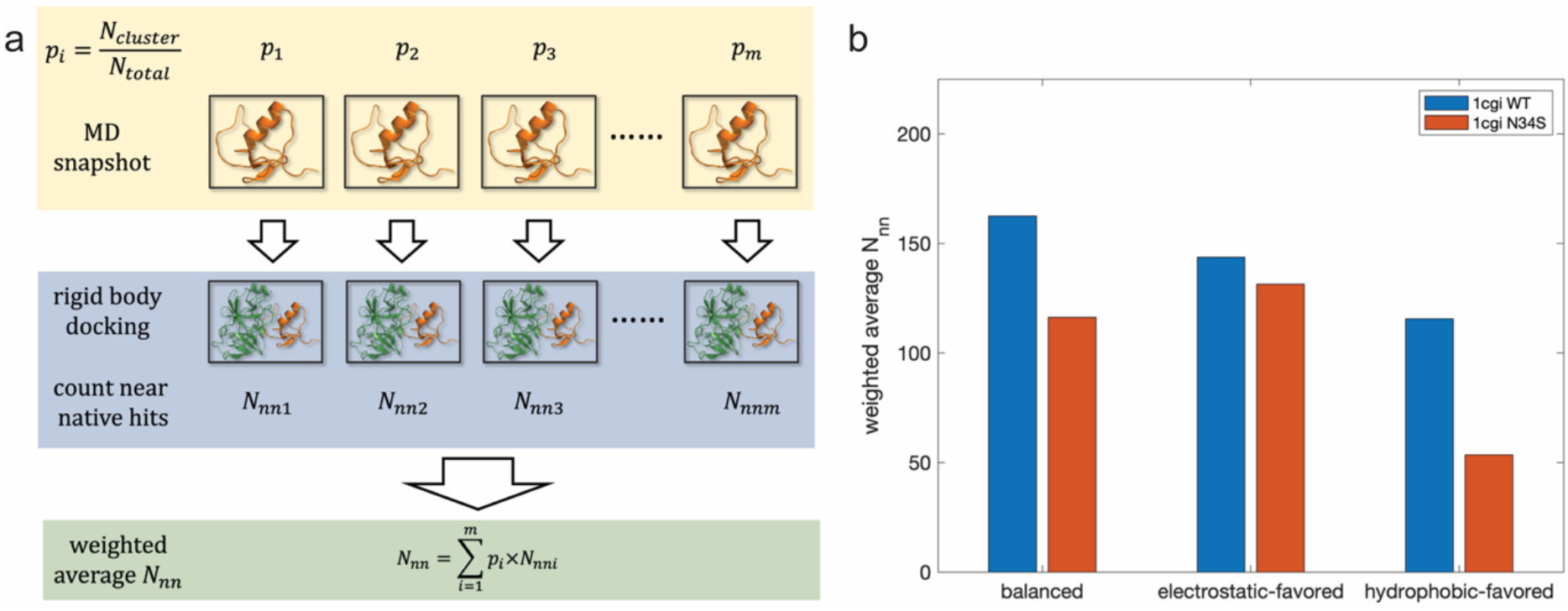
Ensemble docking of wild-type and N34S SPINK1 structures generated by molecular dynamics. **a**. Concept of determining the ensemble average of docked structures by clustering the MD frames from the MD simulations of SPINK1, calculating cluster probabilities, docking the cluster centers, counting the near-native structures from each calculation, and forming the weighted sum to estimate the number of productive encounter complexes. **b**. Weighted average numbers of near-native docked structures from the docking of MD derived cluster centers of wild-type and N34S SPINK1 structures to trypsin (blue and red bars, respectively).

### Ensemble docking of SPINK1 to trypsin reveals reduction in productive collisions

The SPINK1 structures at the centers of the clusters of the MD frames were docked to the trypsin structure (chain A of PDB ID 2ra3) using global protein-protein docking. The trypsin structure is fixed at the center of the coordinate system, and each SPINK1 structure is rotated and translated around it on dense grids. As described in the Methods, we perform 70,000 rotations. At each rotation SPINK1 is translated on a grid with 1 Å step size, and the lowest energy position is retained. After all rotations are completed for the ith cluster, the lowest energy 1000 structures are selected, and the number *N_nni_* of near-native hits is determined. To be considered near-native, a docked pose needed to have interface root mean square deviation (iRMSD) lower than 10 Å from the reference structure discussed in the previous section. We note that iRMSD is defined for atoms that are within 10 Å from any atom of the other protein, so we focus on the interface region rather than of distant parts of the proteins. To account for the effect of docking multiple structures, each representing a cluster with probability *p_i_* and resulting in *N_nni_* docked structures, the weighted average *N_nn_* was calculated for the wild-type and the mutant by *N_nn_* = ∑*p_i_ N_nni_*. Thus, *N_nn_* is obtained by summing the numbers of the near-native docked structures for each cluster, weighted with the probability of the cluster in the MD generated ensemble (**Figure 3a**).

Each docking calculation was carried out by three different sets of parameters. As will be described in Methods, the scoring function evaluating the strength of interactions between two proteins in docking includes four energy terms, the first two representing attractive and repulsive van der Waals contributions describing shape complementarity, the third the electrostatic part of the binding energy, and the fourth the desolvation energy (Kozakov et al., 2017). The relative weighting coefficients of these energy terms for a particular pair of proteins is generally uncertain, and hence we generate three sets of docked structures with parameter sets defined as balanced, electrostatic-favored, and hydrophobicity-favored (Kozakov et al., 2017). **Figure 3b** shows the weighted averages from the docking of the wild-type and N34S SPINK1 to trypsin, and reveals that the mutation reduces the number of docked structures using any of the three energy parameter sets. We note that enzyme-inhibitor complexes are generally stabilized by both electrostatic and hydrophobic contributions to the binding free energy. This is most likely the case for the trypsin-SPINK1 complex, and hence we suggest the use of the balanced set of energy parameters, which in this case shows about 25% reduction in the number of near-native docked structures due to the mutation.

To interpret the docking results we need to briefly discuss the process of protein-protein association. In solution two proteins approach each other due to diffusion. The maximum rate constant, given by the Smoluchowski equation(Berg & von Hippel, 1985; von Hippel & Berg, 1989a), is calculated to be 10^9^–10^10^ M^−1^ s^−1^, on the higher side for charged proteins with strong long-range electrostatic interactions(Alsallaq & Zhou, 2008). However, only a relatively small fraction of collisions results in productive association, and hence one might expect k_assoc_ values around 10^5^ – 10^6^ M^−1^ s^−1^. In fact, the non-specific interactions between two proteins must be somewhat repulsive in order to avoid aggregation(Camacho, Kimura, DeLisi, & Vajda, 2000). The repulsion occurs due to the loss of translational and rotational entropy, and the two molecules stay together only when this loss is compensated by favorable enthalpic or hydrophobic (desolvation) interactions that assume a certain level of shape and electrostatic complementarity. Without such interactions the encounter complex falls apart(Ubbink, 2009). In contrast, if the encounter complex is stable enough, the interface regions will adjust to each other, and the complex moves toward the native state in the conformational space, reducing its binding free energy. The encounter complex can move toward the native state only if it is located in a broad energy well, frequently termed as a “funnel”, which provides sufficiently long initial binding (Camacho et al., 1999; Harel, Spaar, & Schreiber, 2009; Kozakov et al., 2014).

Since productive association between two proteins can occur only from encounter complexes that are sufficiently close to the native state, our hypothesis is that the fraction of docked structures in the near-native region is indicative of the association rate. Although for trypsin and the pancreatic trypsin inhibitor the second-order rate constant for the association is very high, about 1.1 × 10^6^ M^−1^ s^−1^ (Vincent & Lazdunski, 1972), it is still about three orders of magnitude lower than the Smoluchowski limit. Since the docking generates 70,000 structures (one translation for each rotation, see Methods), the 150 structures within the 10 Å RMSD neighborhood of the native structure (see **Figure 3b**) shows a productive ratio of about 0.002, suggesting that selecting this region to define productive encounter complexes is about right. The main conclusion from our simulations is that the N34S mutation reduces the fraction of productive conformations and hence the trypsin-SPINK1 association rate by 10% to 20%. It is important to note that the association between trypsin and SPINK1 is unusually strong. The first-order rate constant for the dissociation is 6.6 × 10^−8^ s^−1^, which corresponds to a half-life of about 17 weeks (Abita, Lazdunski, Bonsen, Pieterson, & de Haas, 1972; Vincent & Lazdunski, 1972). As swill be shown, the 10% increase in the association rate is difficult to measure. However, we will also show that the 10% decrease in the trypsin-SPINK1 association rate has a large impact on the kinetics of trypsinogen to trypsin transition due to the autocatalytic character of the process and the initially low trypsin concentration, which makes the system very sensitive to minor changes in the rate of inhibition.

### Kinetic modeling shows earlier increase of trypsin concentration

Trypsin is a proteolytic enzyme, which is biosynthesized and secreted as the inactive precursor trypsinogen. Such precursors to active enzymes are generally known as proenzymes or zymogens. Trypsinogen becomes activated by limited proteolysis of its 8 amino acid-long N-terminal activation peptide. Ectopic activation of trypsin inside the pancreas occurs through trypsin-mediated trypsinogen activation, commonly referred to as autoactivation(Garcia-Moreno, Havsteen, Varon, & Rix-Matzen, 1991; Geisz & Sahin-Toth, 2018; Wu, Wu, & Wang, 2001). The newly released trypsin in the pancreas is inhibited by SPINK1. Levels in the pancreatic juice amount to about 0.1–0.8% of the total protein(Pubols, Bartelt, & Greene, 1974), which, assuming that about 25% of the juice proteins is trypsinogen and after correction for the molecular mass difference, should translate to SPINK1 concentrations that can inhibit about 13% to 20% of the potential trypsin content(Hegyi & Sahin-Toth, 2017). During autoactivation of trypsinogen, the newly generated trypsin reacts with SPINK1 and becomes unavailable to catalyze further trypsinogen activation. In time, however, SPINK1 reserves become depleted and autoactivation can freely proceed. Thus, the protective role of SPINK1 is to delay trypsinogen autoactivation(Hegyi & Sahin-Toth, 2017; Pfutzer et al., 2000; Witt et al., 2000).

The mechanism of the autocatalytic trypsin activation and of its inhibition is generally written as

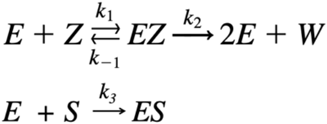

where E and Z represent the enzyme (trypsin) and zymogen (trypsinogen), EZ is the transitional trypsin-trypsinogen complex, W is the peptide which is removed from the trypsinogen, and S is the inhibitor SPINK1 (Garcia-Moreno et al., 1991). In Methods we provide a set of ordinary differential equations to model the kinetics of the above system both with the wild-type and the N34S mutant SPINK1 by assuming a plausible set of rate constants and initial compound concentrations. In **Figure 4a** we present trypsin concentrations with the initial conditions 50 nM and 10 nM for trypsinogen and SPINK1, respectively, thus the amount of SPINK1 is sufficient to inhibit only 20% of the potential trypsin content (Hegyi & Sahin-Toth, 2017). Since trypsinogen always has some trypsin activity, with the trypsin to trypsinogen concentration ratio of 10^−8^, the initial trypsin concentration is assumed to be 0.5X10^−6^ nM. The association rate k3 between trypsin and the native SPINK1 is set as 1.1×10^6^ M^−1^s^−1^ (Vincent & Lazdunski, 1972), resulting in very low trypsin release until about 30 minutes, and then a steep increase to the maximum of 40 nM (blue curve). The red and yellow curves in the same figure show trypsin concentrations if the association rate is reduced by 10% and 20%, respectively, and show that the moderate reduction in the rate leads to early trypsin release that may contribute to chronic pancreatitis due to the digestion of pancreatic tissues. Figure 4a could suggest that the reduced association rate has no impact before about 25 minutes, and then leads to sudden change. However, the inset representing the early stage of trypsinogen activation emphasizes that the three curves start to diverge very early, essentially at the very beginning of the process.

**Figure 4.**
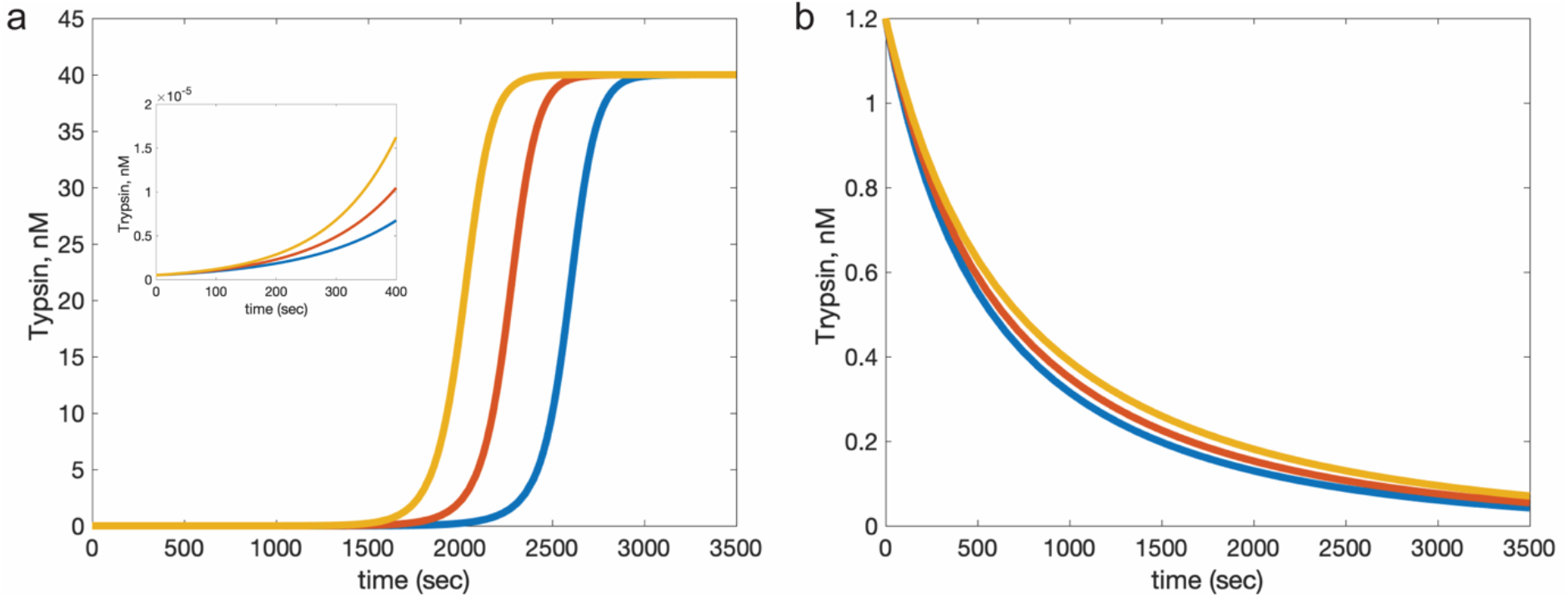
Impact of reduced association rate on trypsinogen activation and trypsin inhibition. **a**. Trypsin concentrations obtained by trypsinogen activation in the presence of wild-type SPINK1 (blue curve) and with N34S SPINK1 assuming the reduction of association rate by 10% (red curve) and by 20% (yellow curve). The inset shows that the three curves start to diverge very early, essentially at time = 0s. **b**. Trypsin concentration in typical inhibition experiments in the presence of wild-type SPINK1 (blue curve) and assuming the reduction of association rate by 10% (red curve) and by 20% (yellow curve).

The main question that remains to be answered is why no experiment detects the impact of the N34S mutation on the association rate. In fact, the accepted experimental observation is that the N34S mutation does not change either the binding affinity or the association rate. Figure 4b show simulation results for a typical binding experiment used to determine the association rate(Vincent & Lazdunski, 1972), and sheds some light on this controversy. The initial conditions are 1.2 nM and 1.8 nM for trypsin and SPINK1, and the blue curve has been generated with association and dissociation rates, respectively, of 1.1×10^6^ M^−1^s^−1^ and 6.6 × 10^−8^ s^−1^. The association rate is then reduced by 10% and 20% (red and yellow curves), while the dissociation rate is not changed. As shown, the impact of the change in the rate constant is very small, in contrast to the large change in the time of trypsin release in the previous simulation. Thus, we suggest that the impact of the N34S mutation can be detected only when measuring the rate of trypsinogen to trypsin conversion in the presence of a less than stoichiometric amount of inhibitor and very low initial trypsin concentration. At the beginning of this process the system is very sensitive to minor changes in the rate of inhibition due to the autocatalytic mechanism of the conversion. A small fraction (on the order of 10^−8^) of trypsinogen always converts to trypsin along a non-autocatalytic pathway (i.e. intrinsic zymogen activity), and hence available to start the catalytic reaction(Pasternak, Liu, Lin, & Hedstrom, 1998). However, removing any fraction of the small amount of trypsin at the early stage of the process by SPINK1 will substantially delay trypsin production. Thus, even a small decrease in the inhibition rate will have a big impact on trypsin release rate as demonstrated in Figure 4a. As shown in Figure 4b, the same change in the association rate is almost undetectable when simply adding inhibitor to trypsin.

## Discussion

We introduced here a new approach to examining the potential pathogenic mechanism of the N34S mutation of SPINK1 that is a leading cause of chronic pancreatitis. We proceeded in three steps. First, we used molecular dynamics to generate ensembles of conformation for wild-type and N34S SPINK1, and showed that the mutation slightly rigidifies the loop region containing residue 34. Second, we docked SPINK1 structures representing the ensemble to a structure of trypsin, and showed that the N34S mutation reduces the number of productive encounter complexes and hence the rate of SPINK1-trypsin association. Third, kinetic modeling revealed that even a 10% change in SPINK1-trypsin association rate had major impact on the rate on trypsinogen activation and thus trypsin release. The main argument is that since the initial trypsin concentration is very low, it is substantially affected by any change in the association rate with the inhibitor, and the effect is many times amplified by the autocatalytic mechanism of trypsinogen activation that makes trypsin release rate highly sensitive to initial trypsin concentration, which in turn is affected by the inhibition rate. We also modeled experiments of mixing native and N34S SPINK1 with trypsin to determine the association rate, and has shown that under the typical experimental conditions the impact of 10% and even 20% change in the association rate has very small impact on the time course of trypsin concentration, well within the error bar of detection methods.

Our analysis suggests that the best approach to validate the above mechanism would be to perform trypsinogen activation experiments(Geisz & Sahin-Toth, 2018) in the presence of inhibitors. In these experiments we consider some amount of trypsinogen, add sub-stoichiometric amounts (say, 20%) of SPINK1, add a very small amount of trypsin, and measure the concentration of trypsin as the function of time. The same experiments should be performed both with wild-type and N34S SPINK1. However, such experiments failed in several laboratories, including ours. The problem is that a small fraction (on the order of 10^−8^) of trypsinogen converts to trypsin in a non-trypsin catalyzed process, and hence trypsinogen always contains trace elements of trypsin(Pasternak et al., 1998). Since the rate of trypsinogen activation is extremely sensitive to the initial trypsin concentration, which is difficult to control, this type of experiments provide almost no information due to the limited reproducibility of the results. Thus, the difficulty of performing experiments increases the importance of the computational approach considered in this paper.

## Methods

### Molecular dynamics simulations and clustering of frames

The conformational changes of the wild-type SPINK1, extracted from the co-crystalized form (PDB ID 1cgi), and the N34S mutant were studied by molecular dynamics (MD) simulations. We used the AMBER99SB-ILDN (Lindorff-Larsen et al., 2010) force field and the TIP3P water model. Before setting up systems and MD parameters, all proteins were processed by Maestro GUI’s Protein Preparation Wizard (Sastry, Adzhigirey, Day, Annabhimoju, & Sherman, 2013). The simulations were performed using the GPU version of Desmond (Bowers et al., 2006) on a desktop computer with four Nvidia GTX 1080 graphics cards. Before production runs, every simulation started with the built-in standard Desmond equilibration and relaxation protocol. The production runs were configured NPT (T = 310 K) using Nose–Hoover chain with a 1 ps relaxation time for thermostat (single temperature group), and Martyna–Tobias–Klein barostat with 2 ps relaxation time and isotropic coupling. The multiple time step RESPA integrator was set to Δt= 2.5 fs for bonded and near non-bonded interactions and Δt = 7.5 fs for far nonbonded, a previously used standard protocol (Ignatov et al., 2019). Water molecules were constrained with SHAKE. Each simulation was set to run for 500 ns, with recording intervals of 1000 ps. For each protein, we ran 10 independent simulations each starting with random initial velocities drawn from a Gaussian distribution corresponding to atomic kinetic energies that add up to T = 310 K by the equipartition principle, as an effort to average the noisiness of MD simulations. The resulting 10 trajectories were concatenated to represent an aggregate of 5 μs MD simulation. For RMSD analysis of the trajectories, we used the Desmond utility program Simulation Event Analysis implemented in Maestro (Schrödinger, 2017). The RMSD values were calculated referencing the bound conformation of SPINK1 in the PDB structure 1cgi based on all Ca atoms.

To extract representative structures from the MD simulations, we subjected the trajectories to clustering. First, using the Desmond Trajectory Clustering program (Bowers et al., 2006) implemented in Maestro(Schrödinger, 2017), pairwise fitted interface root-mean-square deviation (RMSD) matrices for all frames were generated based on the Ca atoms of the proteins. A greedy clustering algorithm, which finds nearest neighbors within a certain radius, uses those RMSD matrices as the distance measure. The clustering radii were tailored to individual trajectory with respect to pairwise RMSD distributions, based on our previous experience (Ignatov et al., 2019; Kozakov et al., 2005). The general thought process was the following: first, if there was a bimodal distribution, the minimum between the two peaks was chosen as the optimal clustering radius; if there was no such distribution, the peak RMSD value of the distribution was treated as an optimal clustering radius. The clustering radii we eventually applied were 2.8 Å and 2.6 Å for the wild-type and the N34S mutant, respectively. Finally, the cluster centers, which were structures with the largest numbers of neighbors, were extracted from the trajectories using Desmond utilities (Bowers et al., 2006). Only “significant” cluster centers were saved for the next step. More specifically, if a cluster had lower than 2% of total population, meaning that the cluster consisted of fewer than 100 frames out of the total 5020 frames from the concatenated trajectories, that cluster was assessed to be “insignificant”. Based on these criteria, 12 wild-type cluster centers and ten N34S mutant cluster centers were extracted as representative structures from the simulations, each with its probability *p_i_* as described earlier.

### Protein-protein docking

The centers of the clusters of MD frames were docked to the trypsin structure (chain A of PDB ID 2ra3), using the docking program PIPER, which performs rigid body docking in the 6D space of rotations and translations using the fast Fourier transform (FFT) correlation approach. The center of mass of trypsin was fixed at the origin of the coordinate system, and the possible rotational and translational positions of the SPINK1 structure were evaluated at the given level of discretization. The rotational space was sampled on a sphere-based grid that defines a subdivision of a spherical surface in which each pixel covers the same surface area as every other pixel (Yershova, Jain, LaValle, & Mitchell, 2010). The 70,000 rotations we considered correspond to about 5 degrees in terms of the Euler angles. The step size of the translational grid was 1 Å, and hence the program evaluated the energy for 10^9^-10^10^ conformations. For each rotation we retained the translation with the lowest energy value, and thus generated 70,000 docked structures. After performing all rotations, only the 1000 lowest energy structures were retained for further analysis.

The energy function used for docking was a linear combination of the repulsive and attractive contributions to the van der Waals interaction energy, an electrostatic energy term, and a desolvation term, represented by a pairwise structure-based potential based on the Decoys as the Reference State (DARS) (Chuang, Kozakov, Brenke, Comeau, & Vajda, 2008) approach. Since the importance of different contributions to the energy function is somewhat uncertain, we generated three sets of models using the scoring schemes called balanced, electrostatics-favored, and hydrophobicity-favored (Kozakov et al., 2017). In the balanced set the weighting coefficients are selected to yield similar weights for the four different energy terms. The set was shown to generally provide good results for enzyme-inhibitor complexes. In the electrostatic-favored set the weight of the electrostatics was doubled relative to the balanced energy expression, and in the hydrophobicity-favored potential we doubled the weight of the desolvation term.

Docking was performed for individual cluster centers from the MD simulations, resulting in 1000 docked structures for each. We then counted the number of near-native hits (*N_nni_*) in these 1000 poses. To be considered near-native, a docked pose needed to have interface root-meansquare deviation (iRMSD) lower than 10 Å compared to the reference structure, and obtained the weighted average *N_nn_* = ∑*p_i_ N_nni_* as shown in Figure 4a.

### Kinetic modeling of trypsinogen activation

The chemical compounds in the simplified kinetic model are trypsin (E), trypsinogen (Z), the inhibitor SPINK1 (S), and the transitional trypsin-trypsinogen complex (EZ). The trypsin-SPINK complex (ES) and the peptide (W) are end products and do not affect the kinetics of the process. For the concentrations of the other four compounds we introduce the notation y_1_ = [E], y_2_ = [Z], y_3_ = [S], and y_4_ = [EZ]. The mass-action type kinetic model is given by

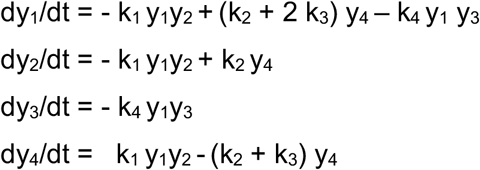

Analogously to the derivation of the Michaelis-Menten equation we assume that the concentration of the enzyme-zymogen complex is small and nearly constant after a brief transitional period and hence use the quasi-steady-state assumption dy_4_/dt = 0, which yields the expression y_4_ = k_1_ y_1_y_2_ /(k_2_ + k_3_), and reduces the above system to three differential equations. The following values were selected for the rate constants: k_1_ = 0.7×10^6^ M^−1^s^−1^, k_2_ = 10^6^ M^−1^, k_3_ = 10^6^ M^−1^, and k_4_ = 1.1 *10^6^ M^−1^s^−1^. As already specified, the ODE’s were solved for the initial conditions y_1_^0^ = 0.5×10^−6^ nM, y_2_^0^ = 50 nM, and y_3_^0^ = 10 nM, thus the amount of SPINK1 was sufficient to inhibit only 20% of the trypsin to be released, and the initial trypsin concentration was small as determined by the trypsin activity of trypsinogen. The differential equations were solved using the ODE solver ode45 ODE solver in MATLAB R2020a.

## Acknowledgments

This investigation was supported by grants DBI 1759277 and AF 1645512 from the National Science Foundation, and R35GM118078, R21GM127952, and RM1135136 from the National Institute of General Medical Sciences.

